# Global and cross-omics feature aggregation improves single-cell multi-omics integration and clustering

**DOI:** 10.1101/2025.04.10.648152

**Authors:** Yingjie Guo, Chenyang Cheng, Zheqing Zhu, Jin Zhou, Haiping Lu, Yuhua Qian, Zhen Liang

**Affiliations:** Institute of Big Data Science and Industry, Shanxi University; School of Computer and Information Technology, Shanxi University; Centre for Machine Intelligence, University of Sheffield; School of Computer Science, University of Sheffield; School of Life Science, Shanxi University

**Keywords:** single cell, multi-omics, data integration, feature aggregation, clustering

## Abstract

Sequencing technology advancements enable the simultaneous measurement of multiple omics within single cells, thereby facilitating single-cell multi-omics analysis. Most existing deep multi-omics integration methods learn shared or omics-specific representations from multiple omics by omics-wise aggregation. However, these approaches often fail to account for the structural relationships among all cells. Here, we introduce scCODI, an innovative cross-omics deep learning model to integrate single-cell multi-omics data for cell clustering. This model processes paired omics datasets, combining single-cell transcriptomics with either epigenomics or proteomics data to facilitate a more comprehensive exploration of cellular heterogeneity. Specifically, the shared representation of two omics is derived through cross-omics feature aggregation, fully leveraging the complementarity of different omics within the same cell. Additionally, we align the omic-specific and shared representations of each cell using a global relationship-guided contrastive learning module, which enhances the similarity between the shared and omic-specific representations of the same cell. Extensive experiments demonstrate that scCODI outperforms previous methods in single-cell multi-omics data integration and cell clustering, as evidenced by its performance across nine paired multi-omics datasets. This makes it a promising tool for decoding complex biological mechanisms. The code is available at https://github.com/SXUI-BDSI/scCODI

## 1. Introduction

Recent advancements in single-cell multi-omics sequencing technologies have facilitated the simultaneous and detailed profiling of multiple molecular modalities, such as RNA expression, protein abundance (ADT), and chromatin accessibility (ATAC) within individual cells [1]. SNARE-seq [2] and SMAGE-seq [3] integrate single-cell RNA data with chromatin accessibility information, and CITE-seq [4–6] links RNA data with protein measurements. These technologies reveal cellular heterogeneity across multiple molecular dimensions and allow more detailed characterization of cellular features to elucidate interactions between distinct omics layers[7]. They provide a holistic view of cellular interactions and regulatory networks at single-cell resolution [8]. Given the intrinsic complementarity of different omics [3], increasing the number of measured modalities further enhances our ability to decode complex biological mechanisms such as cellular function, tissue development, and disease pathogenesis [9]. However, with the rapid expansion of single-cell multi-omics datasets, there is an urgent need for advanced computational frameworks capable of harmonizing high-dimensional data from multiple sources. Such integrative tools are critical to enabling downstream analyses such as clustering and cell-type identification, and unlocking the full potential of increasingly powerful experimental platforms [10].

Cluster analysis plays a pivotal role in single-cell research, and it serves as a foundational method for investigating cellular heterogeneity and biological processes at the level of cell types or subtypes. It provides critical insights that are often obscured in population-level analyses [11, 12]. Commonly used clustering methods for scRNA-seq data include TSCAN and SC3. TSCAN applies principal component analysis (PCA) to reduce the dimensionality of scRNA-seq data, followed by clustering in low-dimensional space using a Gaussian mixture model (GMM) [13]. SC3 employs spectral clustering to generate clustering results from distance matrices based on Euclidean, Pearson, or Spearman metrics. These results are integrated into a shared feature matrix, which is analyzed subsequently through hierarchical clustering to determine the final clusters [14]. Although this is effective for single-omics data, these traditional clustering approaches are not optimized to exploit the complexity of multi-omics datasets, which limits their utility in integrative multi-omics analyses.

Deep learning methods have emerged as a dominant approach for single-cell data analysis because of their exceptional feature extraction capabilities [15]. Recently, various deep learning models have been developed to integrate bimodal data, which advances the integration of multi-omics datasets. For instance, the single-cell Multimodal Variational AutoEncoder (scMVAE) [16] has been utilized to integrate scRNA-seq and scATAC-seq data. This model combines probabilistic Gaussian mixture models with three distinct joint learning strategies to identify latent features that effectively represent multiomics data. However, embedding different omics modalities into a shared latent space can lead to the loss of omic-specific information, which potentially limits the model’s capacity to capture unique omics characteristics.

In contrast to scMVAE, the Deep Cross-omics Cycle Attention (DCCA) model employs deep generative networks to process scRNA-seq and scATAC-seq data individually. It incorporates attention-transfer techniques to model interactions between omics layers while accounting for cellular heterogeneity [17]. scMCs are designed to tackle the individuality in data representation by capturing omic-specific features effectively, enabling better characterization of omic-specific variations. However, their strong emphasis on omic-specific features often limits the ability to extract shared characteristics across different omics layers [18]. The scMDC model integrates RNA and ATAC data or RNA and ADT data by directly concatenating the raw data and using an encoder to derive latent feature representations suitable for clustering [19]. scMDC employs a Zero-Inflated Negative Binomial (ZINB) model [20, 21] to represent single-cell RNA data more accurately, and it addresses dropout events that methods such as BREM-SC [22] and TotalVI [23] have limitations in to account for during the embedding or clustering process. However, scMDC also faces challenges to capture specific omics representations adequately.

This paper presents a novel single-cell multi-omics integration method, scCODI (single-cell Cross-Omics Data Integration), designed to integrate and analyze multimodal single-cell data. The scCODI workflow, illustrated in Figure 1, begins with the preprocessing of paired RNA and ATAC (or ADT) data from SNARE-seq, SMAGE-seq, and CITE-seq. After preprocessing, the data from two modality RNA and ATAC (or ADT) are concatenated to generate an integrated dataset. Both the integrated and single modal datasets are then fed into scCODI, where they undergo processing through two key modules: the global and cross-omics feature integration (COFInt) module and the global relationship-guided contrastive learning (GRgCL) module. The COFInt module performs global and cross-omics feature aggregation to obtain a shared representation of RNA and ATAC (or ADT) data, fully exploiting the complementarity between different omics within the same cell. The GRgCL module employs contrastive learning to align shared and omic-specific representations, ensuring the representations of the same cell across different omics become more similar. Additionally, it effectively mitigates the issue of negative pairs from the same cluster but belonging to different cells, which would otherwise result from low similarity scores. Together, these modules produce embeddings suitable for clustering. The resulting clusters are subsequently used for downstream tasks such as marker detection and pathway analysis. Extensive evaluations on SNARE-seq, SMAGE-seq, and CITE-seq data demonstrate that scCODI significantly outperforms previous methods in clustering multimodal single-cell data.

**Figure 1.**
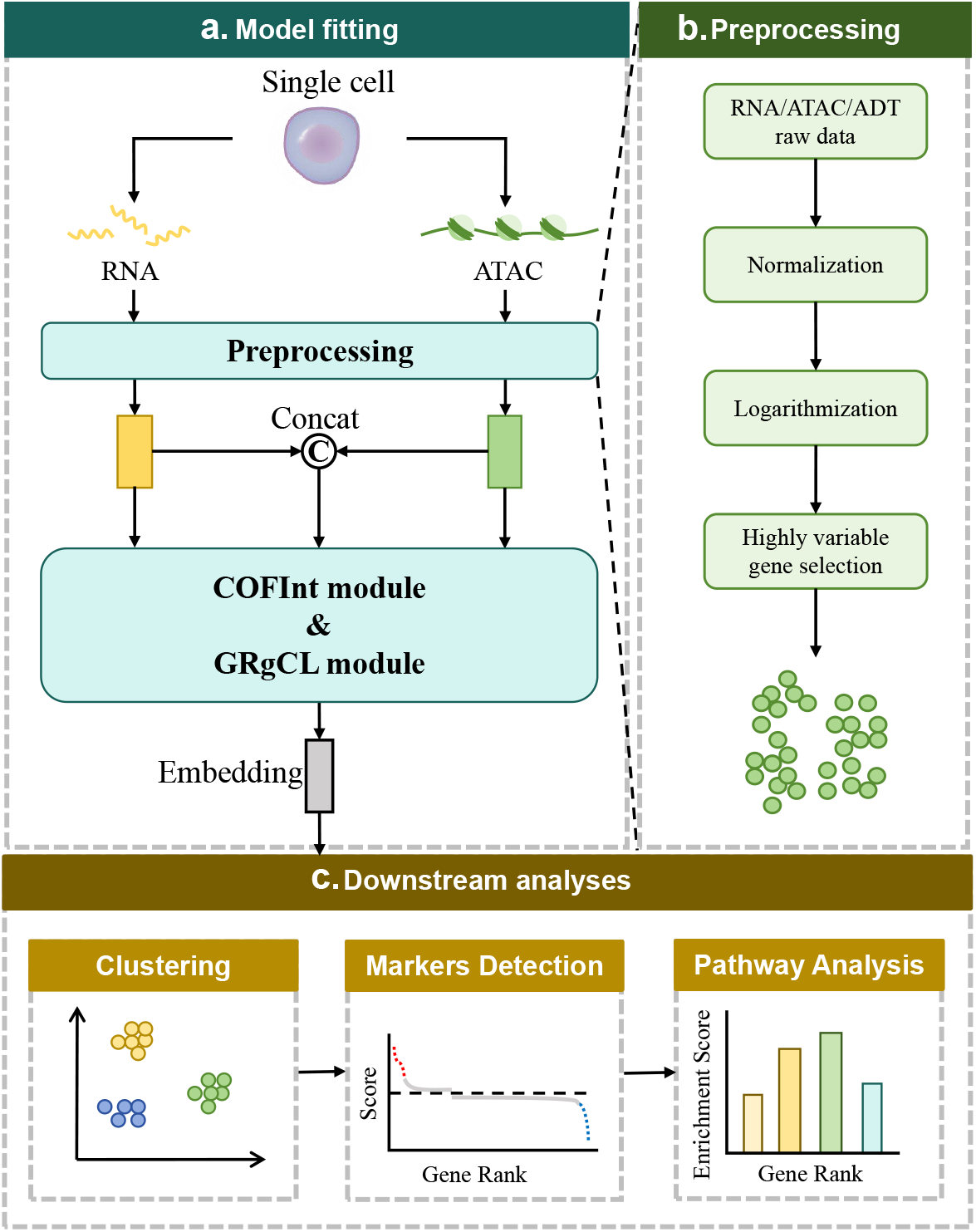
Overview of the scCODI workflow. The workflow begins with model fitting (a). Using single-cell multi-omics technology, RNA and ATAC (or ADT), can be obtained from the same cell. These raw data then undergo preprocessing (b), which includes normalization, logarithmic transformation, and the selection of highly variable genes to reduce noise and prepare the data for integration. By concatenating the preprocessed RNA and ATAC, an integrated input is obtained. This integrated input and preprocessed RNA and ATAC, are separately fed into the the Cross-Omics Feature Integration (COFInt) module for further data augmentation. Subsequently, the global relationship-guided Contrastive Learning (GRgCL) module employs contrastive learning to align shared and omic-specific representations. It selects negative samples using the global relationship matrix obtained from COFInt and treats the same cell’s representations in both shared and omic-specific spaces as positive samples, thereby incorporating omic-specific information to generate the final embeddings. These embeddings are then used for cluster analysis and subsequent downstream analysis tasks (c) such as marker gene detection and pathway analysis for deeper biological insights.

## 2. Materials and methods

The network architecture of scCODI (Figure 2) includes three encoders: one for concatenated data and two for single modal data, along with two decoders tailored to each modality, it can be applied to clustering single-cell multi-omics RNA and ATAC (or ADT) datasets. The model integrates two key components: COFInt and GRgCL. COFInt incorporates a transformer structure to capture global relationships across different feature spaces, producing a shared representation based on these relationships. The global relationship-guided contrastive learning module aligns the shared and omic-specific representations by selecting negative samples based on the global relationship matrix provided by COFInt, while treating the representations of the same cell in both shared and omic-specific spaces as positive samples.

**Figure 2.**
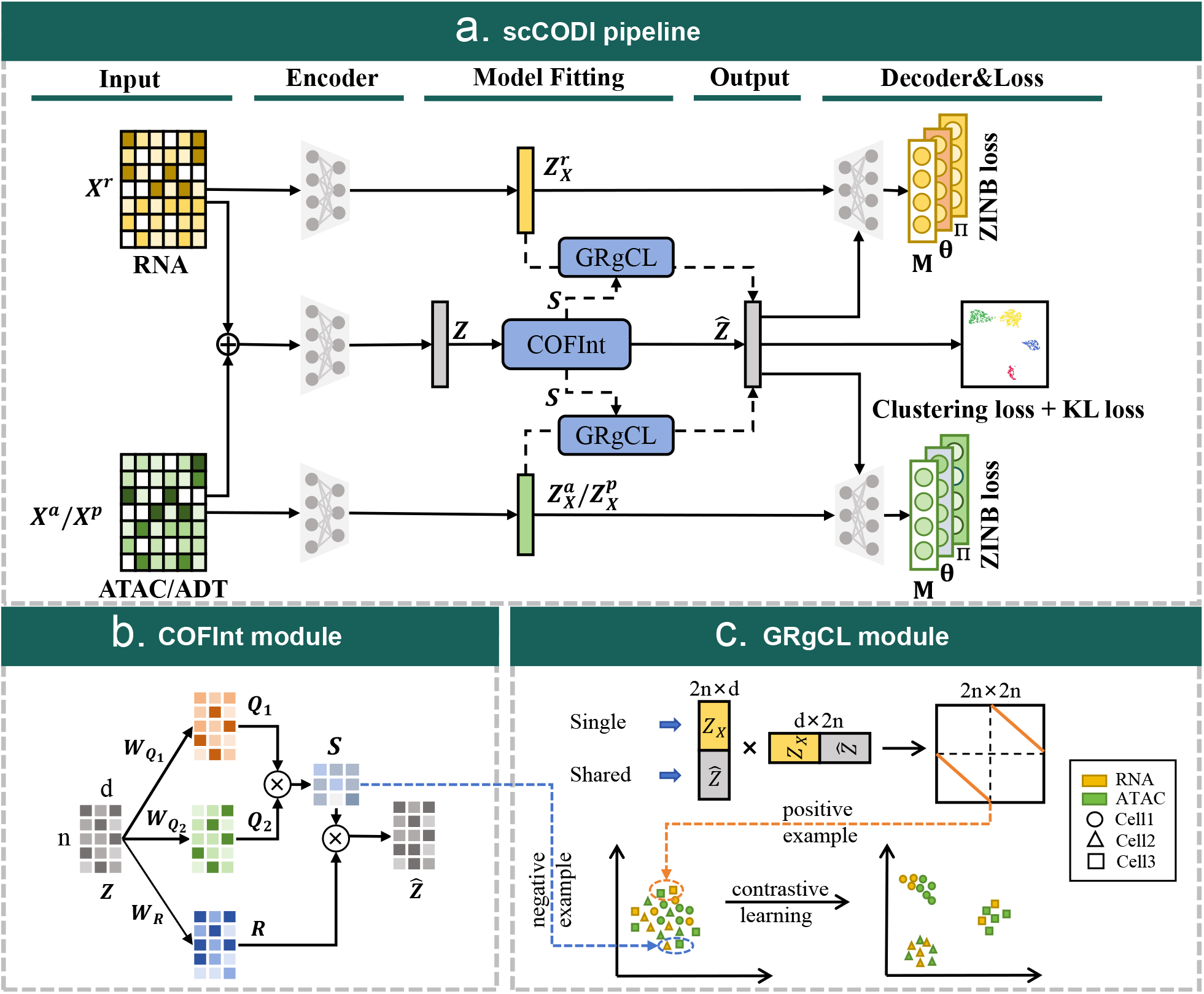
The overall framework and key modules of scCODI. (a) scCODI pipeline: The method integrates multimodal data from RNA and ATAC or RNA and ADT. The model includes an encoder to process concatenated raw data, along with two omic-specific encoders for each omic, and two decoders tailored to each omic. The encoded features are then passed through the Cross-omics Feature Integration (COFInt) module, which performs global and cross-omics feature aggregation, followed by the GRgCL (global relationship-guided Contrastive Learning) module for contrastive learning based on global relationships. The output consists of clustering representations and reconstructed data, optimized through clustering loss, KL divergence loss, and ZINB loss. (b) COFInt Module: The COFInt module utilizes matrices 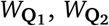, and *W*_**R**_ for feature space transformation across omics, while **S** captures the global relationships among cells. (c) GRgCL Module: The GRgCL module applies contrastive learning to align shared and omic-specific representation. The blue dashed line indicates the selection of negative sample pairs, guided by the global relationship matrix **S**, while the orange dashed line denotes the identification of positive sample pairs, where the same cell exhibits similar representations in both the single modal representation and the shared representation.

### 2.1. Count data preprocessing

The raw data from SNARE-seq and SMAGE-seq are preprocessed and normalized using the Python package SCANPY [24]. The entire preprocessing process is illustrated in (Figure 1b). For each cell, the counts are normalized by a size factor that is denoted as *s*_*i*_, with 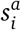 for ATAC data and *s*^*r*^ for RNA data, which is calculated by dividing the library size of the cell by the median library size across all cells. This normalization ensures comparability by equalizing library sizes across cells. Subsequently, the counts are transformed logarithmically and standardized to achieve unit variance and zero mean. The processed count data for RNA and ATAC are utilized in the denoising multimodal autoencoder model, while the raw count matrix is retained for computing the zero-inflated negative binomial (ZINB) loss [15, 25]. CITE-seq data undergo a similar preprocessing and normalization procedure, with the size factor 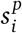 for ADT data calculated in the same manner.

### 2.2. Denoising multi-omics data encoder

In scCODI (Figure 2), RNA data is denoted as 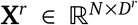, ATAC data as 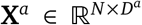, and ADT data as 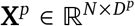. Autoencoders, a fundamental type of neural network, are highly effective for learning nonlinear representations of data [26]. Among their variants, denoising autoencoders are particularly well-suited for handling noisy datasets because they learn robust latent representations by reconstructing clean inputs from corrupted versions [27]. Given the inherently noisy nature of single-cell data, we employed denoising autoencoders for processing RNA, ADT, and ATAC data. The preprocessed RNA, ATAC, and ADT data are denoted as **X**^*r*^, **X**^*a*^, and **X**^*p*^, respectively. After introducing noise, these datasets are represented as 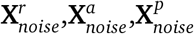, which are formally defined as:

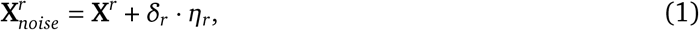

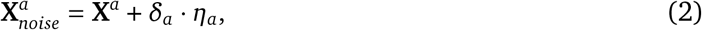

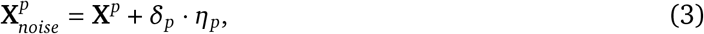

For RNA, ATAC, and ADT data, we add artificial Gaussian noise *η*_*r*_, *η*_*a*_, and *η*_*p*_, respectively. The noise weights *δ*_*r*_, *δ*_*a*_, and *δ*_*p*_ are set to 2.5 for all three data types.

Our autoencoder architecture includes a shared encoder for handling concatenated raw data, two omic-specific encoders for specific omics, and two decoders tailored to different omics datasets. Specifically, for the concatenated data, we denote the encoder for the concatenated RNA and ATAC data, and the encoder for the concatenated RNA and ADT data, as follows:

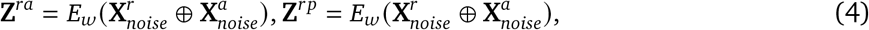

Decoders are defined as:

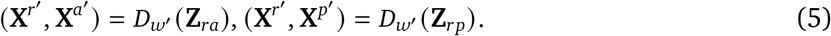

Regarding single omics data, we denote the encoder for the RNA, ATAC, and ADT data as:

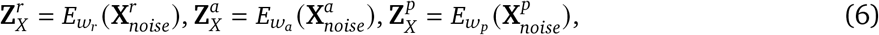

The decoder is defined as:

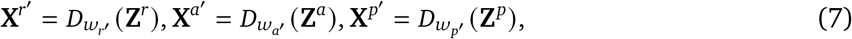

Here, **X**^*r*^′, **X**^*a*^′, **X**^*p*^′, represent the reconstructed data for RNA, ATAC, and ADT, respectively. *w* and *w*′ correspond to the learnable weights of the encoder and decoder. The operator ⊕ denotes the concatenation of two matrices. For all the hidden layers within the encoders and decoders, the Exponential Linear Unit (ELU) [28] activation function was used. Additionally, dropout regularization and batch normalization were applied to the layer output to improve generalization and to stabilize training.

For all omics data, we used zero-inflated negative binomial (ZINB) as the reconstruction loss function [20]. The ZINB model is defined by three key parameters: the mean (*μ*_**X**_) and dispersion (*θ*) parameters of the negative binomial distribution, and a dropout probability coefficient (*π*) that accounts for zero-inflation due to dropout events. Formally, the model can be expressed as:

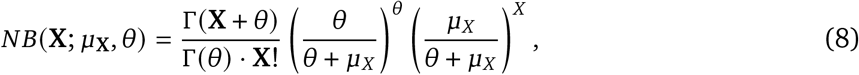

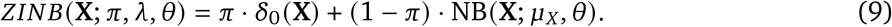

To estimate parameters in the ZINB loss functions, we added three independent fully connected layers **M, Θ**, and **Π** to the last hidden layer of each decoder. The layers are defined as:

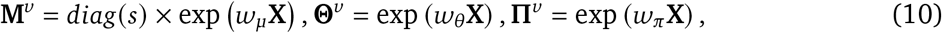

Here, let 𝒟 represent the set of data types, where 𝒟 = RNA, ATAC, ADT. For each data type *v* ∈ 𝒟, we defined the following matrices: **M**^*v*^ as the estimated mean matrix, *θ*^*v*^ as the estimated dispersion matrix, and **Π**^*v*^ as the estimated dropout probability matrix. 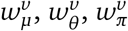 are the learnable weights. The size factors *s*^*v*^ for each data type were determined in the preprocessing step.

Unlike traditional autoencoders that rely on mean squared error loss, the ZINB-based decoder network uses a loss function defined by the negative log-likelihood of the ZINB distribution:

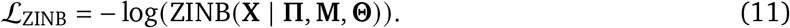

Thus, the reconstruction loss for our autoencoder model using ZINB is the sum of the ZINB loss for specific omics and the ZINB loss for integrated omics data. For the integration of RNA and ATAC, the reconstruction loss is:

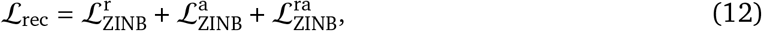

where 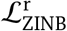 and 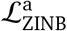 are the ZINB losses for specific RNA and ATAC omics data, respectively, and 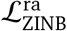 is the ZINB loss for the integrated RNA and ATAC data.

For the integration of RNA and ADT, the reconstruction loss is:

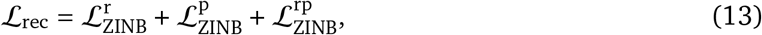

where 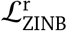 and 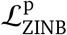 are the ZINB losses for specific RNA and ADT omics data, respectively, and 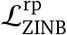 is the ZINB loss for the integrated RNA and ADT data.

### 2.3. Global and Cross-omics Feature Integation

Although concatenating preprocessed data and directly inputting it into the autoencoder can facilitate learning shared features across different omics, this approach may overlook omic-specific patterns, the resulting data representations often contain omic-specific information that may be irrelevant or misleading. This issue can also impact the quality of clustering negatively [29]. Instead of simply using a weighted sum, we enhanced the shared representations of samples across different omics by focusing on those samples with strong inter-relationships within the same cluster.

To obtain a shared representation of multi-omics data, we first modeled the global relationships among samples and utilized these relationships to derive the shared feature representation (Figure 1b). Initially, the raw data was concatenated and input into the autoencoder, which resulted in the shared feature representation **Z**, which is expressed as follows:

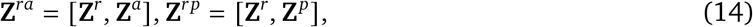

Where 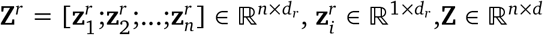. Here, *n* represents the number of samples, and *d*_*r*_ represents the dimensionality of RNA data features in the shared feature representation derived from the autoencoder. The shared feature representations for ATAC and ADT data were obtained using the same method.

Inspired by the transformer attention mechanism [30], we projected **Z** into different feature spaces using **W**_*R*_ to facilitate cross-omics integration across all modalities:

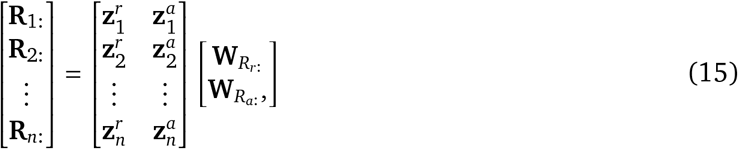

Where 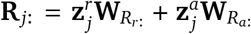; the joint representation mapping of RNA and ADT data can also be obtained using this method. Similarly, **Q**_1_ and **Q**_2_ were obtained using **W**_*Q*1_ and **W**_*Q*2_:

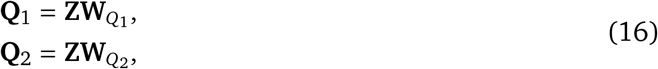

where **Q**_1_, **Q**_2_ ∈ ℝ^*n*×*d*^. The parameters in this module are denoted by **W**_*Z*_ = {**W**_*Q*1_, **W**_*Q*2_, **W**_*R*_}.

The global relationship among samples is represented as follows:

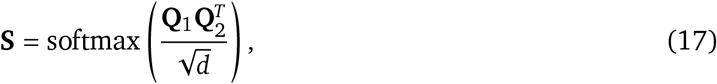

where **S** is a matrix obtained by applying softmax normalization to the inner product of **Q**_1_ and **Q**_2_, which is scaled by 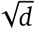.

This matrix **S** enhances the learned representation matrix **R** by incorporating global relationships among samples. Specifically, it allows each sample’s representation to benefit from those of other samples with high correlations, thereby promoting similarity among data representations of samples within the same cluster. The enhanced shared feature representation 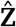 is computed as follows:

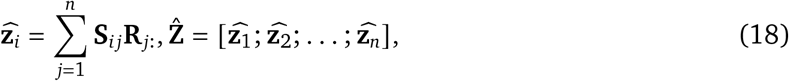

where **R** ∈ ℝ^1×*d*^ represents the representation of the *j*-th sample in matrix **R**, and **S** denotes the relationship weight between the *i*-th and *j*-th samples. The resulting matrix 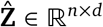.

### 2.4. Global relationship-guided contrastive learning

We enhanced the learning of the common representation 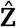 by considering the global relationship matrix across all samples. Because samples within the same cluster should have similar shared feature representations across different omics data, their specific representations within each omics data should also be similar. Therefore, the shared feature representations from the same cluster and the omic-specific representations 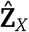 should be mapped close to 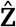 [31, 32]. Based on this, we designed a global relationship-guided contrastive learning method [33], where the shared feature representation and the specific representation of the same sample under different omics were treated as positive pairs. For handling negative pairs, directly setting other samples as negative pairs may lead to inconsistent representations within the same cluster, which contradicts our clustering objective. To address this, we introduced a global relationship-guided multi-omics contrastive learning method (Figure 1c). Specifically, we computed the cosine distance between the shared feature representation 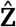 and the specific representation 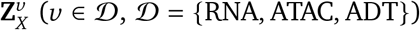 to indicate their similarity:

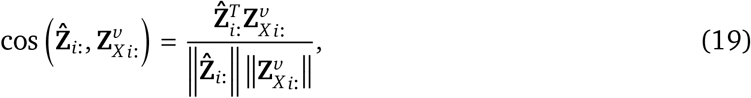

The loss function of the global relationship-guided contrastive learning is defined as:

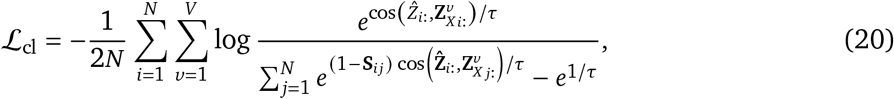

In this equation, the relationship between cos 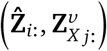 and **S**_*i j*_ is determined by the value of *S*_*i j*_ from Eq. (17) and the temperature parameter *τ*. When *S*_*i j*_ is small, (1 − **S**_*i j*_) is large, which means that the contribution of 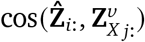 to the loss function is large. Even if 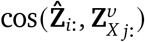 is small, it still occupies a large proportion in the denominator, thereby affecting the overall loss value.

### 2.5. KL divergence in the latent layer

In cluster analysis, it is crucial to group similar data points into the same cluster. We incorporated a Kullback-Leibler (KL) divergence loss function [34], to strengthen the relationships between similar cells while preventing cluster centroids from becoming overly concentrated in the latent space. Similar to t-SNE [35], we utilized a *t*-distribution kernel function to represent the pairwise similarity between two cells *i* and *i*′ in the latent space of our autoencoder:

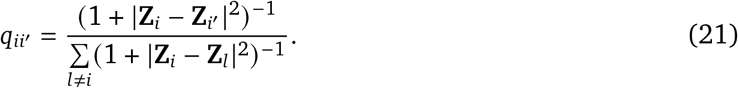

Here, *q*_*ii*_ = 0. During training, the target distribution *P* adjusts affinities between cells by amplifying the similarities of highly similar cells and diminishing those with low similarities. *P* is derived by the square of *Q*, followed by normalization:

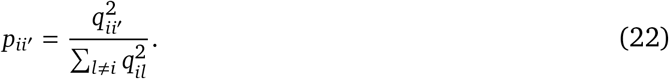

With these two similarity distributions, **Q** and the corresponding target distribution **P**, the KL divergence-based loss function is defined as follows:

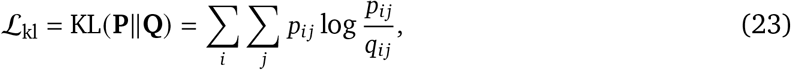

this loss function measures the difference in probability distributions between *P* and *Q*. During training, **P** and **Q** were calculated per batch.

### 2.6. Deep K-means clustering objective function

We apply unsupervised clustering to the latent representations generated by the model [34]. The autoencoder learns a nonlinear mapping for each cell, which transforms the original concatenated data into a low-dimensional space 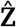 through cross-omics integration. The loss function ℒ_*c*_ for K-means clustering is defined as:

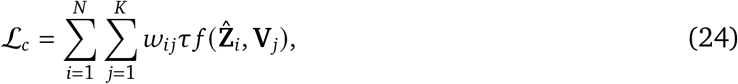

In this equation, **V** represents the *K* clustering centroids, and *f* computes the Euclidean distance between a cell in the latent space and a given centroid. The hyperparameter *τ* is assigned different values depending on the data type. To enhance the smoothness of the gradient descent optimization process, weights were determined using a Gaussian kernel function:

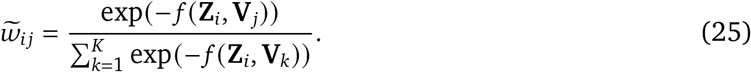

To accelerate convergence, the weights were inflated, and the final *w*_*i j*_ was given by:

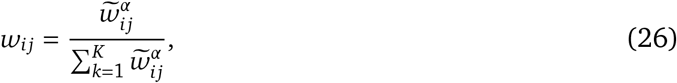

where the hyperparameter *α* is assigned a value of 2.

In the proposed framework, the total loss in our network is defined as:

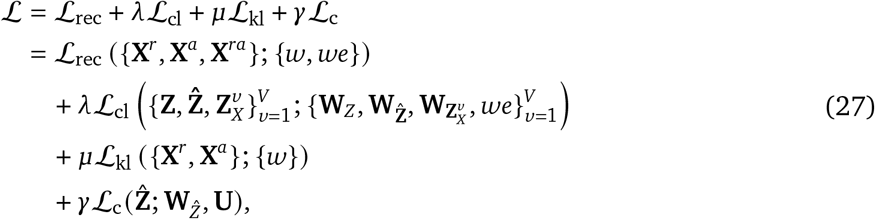

where ℒ_rec_ represents the reconstruction loss of three types of data, which include RNA, ATAC, and the joint reconstruction of the original RNA and ATAC data. The RNA and ADT data are processed similarly. *w* and *w*′ are the parameters of the encoder and decoder. ℒ_cl_ represents the loss induced by contrastive learning, which captures the discrepancy between the relationship-enhanced shared feature representation and the omic-specific representations of specific omics data. ℒ_*c*_ denotes the clustering loss, where the latent representations iteratively converge toward the centroids initialized by K-means, which are represented as *U*. In this framework, *λ, μ*, and *γ* are hyperparameters that regulate the weights of contrastive loss, KL divergence loss, and K-means clustering loss, respectively.

### 3. Results

### 3.1. Experimental setup

#### Datasets

scCODI integrates different sets of single-cell omics data. In our experiments, we evaluated the model’s ability to integrate RNA with chromatin data and RNA with protein data.

For the integration of RNA and ATAC, we used four preprocessed single-cell multi-omics datasets with paired profiles from a previous study [16]: (i) CellMix, which comprised 1,047 cells, obtained from GEO (GSE126074), where gene expression and chromatin accessibility were co-assayed simultaneously in each single cell using SNARE-seq; (ii) AdBrain, which consisted of 10,309 cells, also downloaded from GEO (GSE126074), where gene expression and chromatin accessibility were measured in single cells derived from the adult mouse cerebral cortex; and (iii) Human peripheral blood mononuclear cells (PBMCs) datasets, which consisted of approximately 3,000 cells and 10,000 cells.

For the integration of RNA and ADT, we collected five single-cell multi-omics datasets and preprocessed them: (i) Cord blood mononuclear (BMNC) cells with 1128 cells downloaded from GEO (GSE100866). the data were annotated using marker genes and ADTs [22]; (ii) Bone marrow mononuclear cells with 30,672 cells were downloaded from GEO (GSE128639), and the cell type labels were downloaded from the “bmcite” dataset in the “SeuratData” package (v0.2.1); (iii) The PBMC dataset with 3762 cells that was available on the 10X website. We downloaded the preprocessed data and the cell type labels from the GitHub of Specter [36]; (iv) Mouse spleen lymph node with 16,828 cells was downloaded from GEO (SLN111, GSE150599); and (v) Mouse spleen lymph node with 15,820 cells was downloaded from GEO (SLN206, GSE150599). TotalVI provided cell-type labels on GitHub.

#### Evaluation protocols

For our multi-omics data integration clustering, we applied the K-means algorithm to the final low-dimensional co-embedding representation 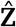. Subsequently, we evaluated the clustering performance using Normalized Mutual Information (NMI) [37] and Adjusted Rand Index (ARI) [38]. Both NMI and ARI values ranged from 0 to 1, with higher values indicating better clustering performance.

#### Comparing methods

We implemented scCODI using the PyTorch deep learning framework and compared its performance with other single-cell multi-omics data integration methods.

For the integration of RNA and ATAC, we used the following methods: (i) **scMCs**, which employed a deep learning model for joint representation of multi-omics data, mapping them to a shared lowdimensional space and enhancing the relationships between different omics through contrastive learning to improve clustering performance [18]; (ii) **scMDC**, which used a multi-branch neural network to learn shared representations and applied clustering algorithms for identification of cell clusters [19]; (iii) **scMVAE**, which incorporated three strategies—scMVAE-PoE, scMVAE-NN, and scMVAE-Direct—for learning joint latent features to facilitate data fusion and clustering. Specifically, scMVAE-Direct concatenated the raw features from each omics dataset, scMVAE-NN integrated lowdimensional features extracted from different omics, and scMVAE-PoE utilized the product of experts framework to estimate a joint posterior distribution [16]; and (iv) **DCCA**, which projected different omics into their respective low-dimensional spaces and used a ‘teacher-student’ mechanism to fuse multi-omics data [17].

To integrate RNA and ADT, we compared with two multi-omics integration methods for RNA and ADT data integration. (i) **BREM-SC** [22] uses a Bayesian model to embed RNA and protein data into a shared low-dimensional space for joint analysis; and (ii) **TotalVI** [23], which is a deep generative model designed specifically for the joint analysis of single-cell RNA and protein data. It utilizes variational inference to model the complex relationships between transcriptomics and proteomics data. Additionally, we used the **scMDC** deep learning method for comparative experiments. scMDC demonstrates robust performance in integrating CITE-seq data, validating its effectiveness across different multi-omics modalities.

### 3.2. Evaluation on real RNA and ATAC integration data

Initially, we evaluated the clustering performance of scCODI on SNARE-seq and SMAGE-seq datasets, benchmarking it against 10 alternative methods. These approaches included models designed specifically for multimodal data clustering (DCCA, scMVAE, scMDC, and scMCs), two single-modal data clustering tools (SC3 and TScan), and two general clustering algorithms (IDEC and K-means). We evaluated these methods using four paired RNA and ATAC datasets; two datasets were derived from SNARE-seq and two were derived from SMAGE-seq (Table 1).

**Table 1.** Performance of clustering for different methods on RNA and ATAC integration datasets. The symbol ‘±’ represents the standard deviation over five repeated experiments. The best results are highlighted in **bold font**, and the second-best results are indicated by underline.

| | GSE126074 | | Adbrain | | SMAGE10× 3k | | SMAGE10× 10k | |
| --- | --- | --- | --- | --- | --- | --- | --- | --- |
|  | NMI | ARI | NMI | ARI | NMI | ARI | NMI | ARI |
| SC3 | <u>.942±.000</u> | <u>.966±.000</u> | .418±.003 | .319±.001 | .551±.003 | .510±.002 | .501±.001 | .446±.002 |
| Kmeans | .726±.000 | .877±.000 | .383±.000 | .143±.000 | .464±.000 | .352±.000 | .501±.000 | .343±.000 |
| Tscan | .772±.001 | .804±.001 | .253±.003 | .189±.003 | .258±.002 | .120±.004 | .401±.001 | .376±.001 |
| IDEC | .518±.001 | .460±.001 | .149±.005 | .092±.003 | .300±.001 | .079±.001 | .253±.001 | .067±.001 |
| DCCA | .898±.000 | .940±.000 | .357±.002 | .406±.003 | .508±.000 | <u>.583±.000</u> | .560±.000 | .600±.000 |
| scMCs | .869±.000 | .907±.000 | .510±.000 | <u>.554±.001</u> | .278±.001 | .121±.000 | .393±.000 | .224±.001 |
| scMDC | .918±.000 | .952±.000 | <u>.588±.001</u> | .434±.001 | <u>.638±.001</u> | .516±.002 | <u>.589±.000</u> | <u>.613±.001</u> |
| scMVAE-POE | .899±.000 | .935±.000 | .457±.003 | .219±.005 | .503±.001 | .285±.001 | .531±.001 | .310±.001 |
| scMVAE-Direct | .892±.000 | .927±.000 | .302±.003 | .137±.001 | .506±.004 | .305±.001 | .540±.001 | .331±.001 |
| scMVAE-NN | .921±.000 | .952±.000 | .468±.003 | .222±.004 | .499±.000 | .265±.000 | .553±.001 | .311±.000 |
| scCODI | **.947±.000** | **.978±.000** | **.595±.005** | **.584±.006** | **.654±.003** | **.606±.006** | **.663±.001** | **.721±.001** |

Using NMI and ARI metrics across different datasets, multimodal methods consistently outperformed single-modal approaches (Figure 3a). scCODI achieved superior performance across both metrics on all four datasets, surpassing other multimodal methods significantly, such as scMVAE-PoE, scMVAE-Direct, and scMVAE-NN (Figure 3a). This highlights the advantages of scCODI. Among these methods, scMVAE-Direct exhibited the weakest performance, likely due to the increased sparsity and complexity that arise from concatenating high-dimensional features. scMVAE-NN outperformed both scMVAE-Direct and scMVAE-PoE by leveraging a more compact feature space for representation. However, scMVAE failed to capture the unique characteristics of multi-omics data in cell clustering adequately. Conversely, scCODI balances the integration of shared features effectively, which is critical for consistent clustering, with the retention of specific features from single-omics views. Although DCCA uses separate neural networks to project multi-omics data into distinct representation spaces, it underperforms compared with scCODI due to its emphasis on specific omics features and insufficient consideration of shared features that are crucial for consistent clustering. In contrast, scCODI extracts both shared and specific features from different omics datasets simultaneously. Additionally, through contrastive learning, scCODI enhances feature similarity across different omics for the same samples, thereby capturing cellular heterogeneity more comprehensively to achieve superior clustering performance.

**Figure 3.**
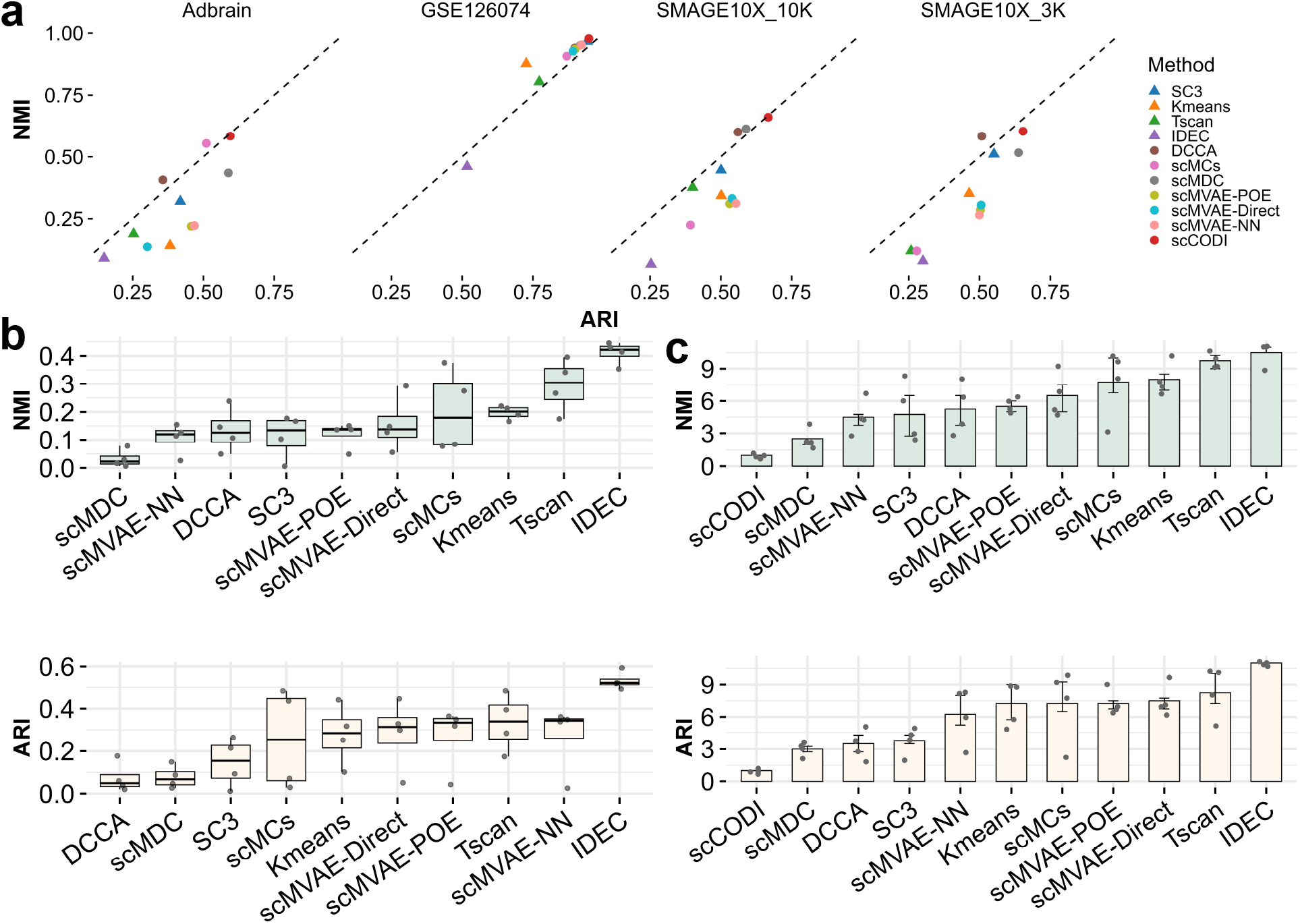
Clustering performance of scCODI and the competing methods on different ATAC datasets. The clustering performance of scCODI and competing methods across different ATAC datasets. All methods were evaluated across four datasets. Panels (a) of the figure visualizes clustering performance in a two-dimensional representation. Panel (b) uses box plots to illustrate the performance differences between scCODI and other methods. Each box plot displays key statistical measures, which include the minimum, first quartile (Q1), median, third quartile (Q3), and maximum values. The interquartile range (IQR), which is the difference between the third quartile (Q3) and the first quartile (Q1). The minimum and maximum are determined as the smallest and largest data points that are greater than or equal to Q1 1.5IQR and less than or equal to Q3 + 1.5IQR, respectively. Each dot represents a data point, which represents a performance difference in a dataset. Panels (c) summarize the average ranking of all methods, with dots indicating a method’s rank for a specific dataset and error bars representing standard errors.

The positive values indicate that scCODI outperforms its counterparts (Figure 3b). The analysis reveals that the median differences in NMI and ARI were approximately 0.1, while the median differences in ARI was around 0.2, which underscores the advantages of scCODI. Furthermore, we ranked all methods based on their performance metrics for each dataset. scCODI consistently achieved the top rank in both metrics after averaging the rank of each method across the four datasets (Figure 3c). The second-best method, scMDC, attained an average rank of 2 for both NMI and ARI. Overall, these results from multiple real datasets demonstrated the stability and robustness of scCODI in clustering performance on SNARE-seq and SMAGE-seq datasets.

Furthermore, to demonstrate the quality of the final low-dimensional co-embedding representation 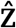, we used uniform manifold approximation and projection (UMAP) [39] to visualize the cell clustering points of scCODI and other baseline methods across various benchmark datasets.The clustering results of baseline methods on AdBrain, SMAGE10X 3K, and SMAGE10X 10K are presented in the Supplementary Material (Figures S3 to S5), which show the visualization and clustering results of different methods on both raw and imputed data. scCODI demonstrated more compact separation of samples within the same cluster and better separation of samples from different clusters (Figure 4). This validated the effectiveness of our proposed data augmentation and contrastive learning methods and explained why scCODI achieved superior clustering performance.

**Figure 4.**
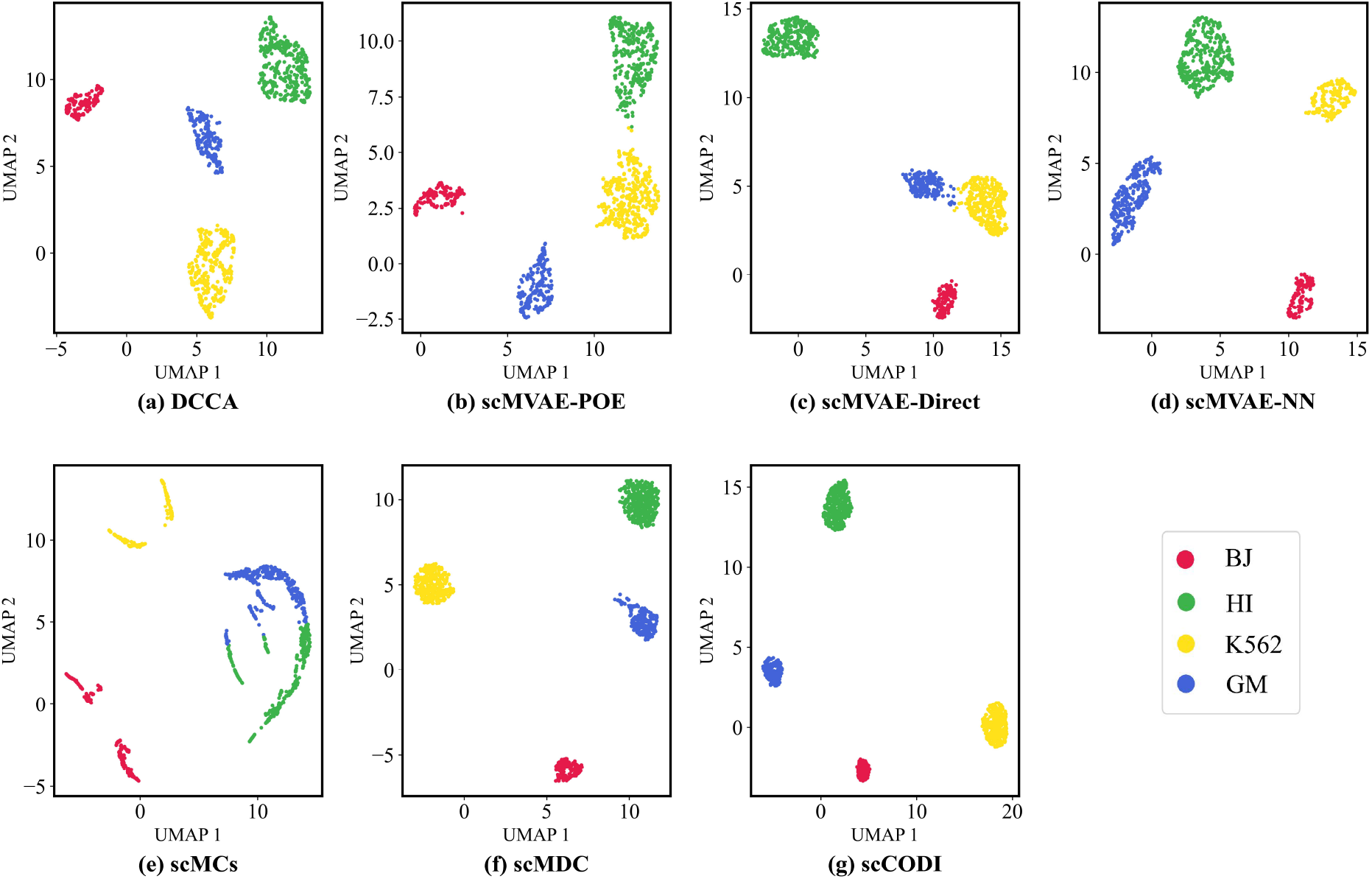
Cell clustering visualization of each method on GSE126074 dataset. (a) DCCA, (b) scMVAE-POE, (c) scMVAE-Direct, (d) scMVAE-NN, (e) scMCs, (f) scMDC, and (g) scCODI.

### 3.3. Evaluation on real RNA and ADT integration data

We evaluated the clustering performance of scCODI on the CITE-seq dataset and compared it with recent competitive methods, which included BREM-SC that was designed for clustering multimodal data, TotalVI, which addresses batch effects, and scMDC, which was developed for learning embeddings of single or multiple modal data. We tested these methods on five CITE-seq datasets. scCODI outperformed the competing methods across all five datasets (Figure S1), except for the PBMCspector dataset, where scMDC exhibited comparable performance. Although TotalVI was less effective than scMDC, it outperformed BREM-SC due to its ability to correct batch effects. The performance differences between scCODI and the competing methods are summarized (Figure S1a). A positive difference indicates superior performance of scCODI. Our analysis reveals that scCODI consistently outperformed all other methods across multiple datasets. Furthermore, we ranked all methods based on their performance metrics for each dataset. scCODI achieved the highest average rank across the five datasets for both metrics consistently (Figure S1c). scCODI ranked first in both metrics across all datasets. In contrast, the second-best method, scMDC, had an average rank of 2 for both NMI and ARI. Overall, our results across various real datasets confirmed that scCODI demonstrated stable and robust clustering performance on CITE-seq datasets (Table 2).

**Table 2.** Performance of clustering for different methods on CITE-seq datasets. The best results are highlighted in **bold font**, and the second-best results are indicated by underline.

|  | BMNC100866 |  | BMNC128639 |  | PBMC_spector |  | spleen111 |  | SLN206 |  |
| --- | --- | --- | --- | --- | --- | --- | --- | --- | --- | --- |
|  | NMI | ARI | NMI | ARI | NMI | ARI | NMI | ARI | NMI | ARI |
| BREM-SC | 0.8036 | 0.7424 | 0.6194 | 0.7285 | 0.5295 | 0.5111 | 0.3991 | 0.2538 | 0.4069 | 0.2765 |
| TotalVI | 0.7876 | 0.8279 | 0.7403 | 0.4645 | <u>0.6455</u> | 0.5016 | 0.6069 | 0.2551 | <u>0.6114</u> | 0.2529 |
| scMDC | <u>0.9679</u> | <u>0.9881</u> | <u>0.8567</u> | <u>0.8934</u> | 0.6377 | <u>0.5160</u> | <u>0.6593</u> | <u>0.5452</u> | 0.6068 | 0.3590 |
| scCODI | **0.9904** | **0.9950** | **0.8631** | **0.9075** | **0.7180** | **0.6149** | **0.6677** | **0.5934** | **0.6281** | **0.4249** |

We used UMAP to visualize the cell clustering of scCODI and other baseline methods across various benchmark datasets after processing the CITE-seq data with our model. scCODI achieved clearer cluster boundaries, lower error rates, and a higher concentration of points within the same cluster (Figure S2). The clustering results on other CITE-seq integration data are presented in the Supplementary Material (Figures S6 to S9). These observations validated the superior clustering performance of scCODI and underscored the effectiveness of our innovative approach.

### 3.4. Evaluation of multimodal integrated data

As discussed in the introduction, different omics data provide unique and complementary information, which is essential for accurate cell clustering and identification of cell types. Consequently, integrating multi-omics data is expected to yield superior clustering performance compared with approaches that rely solely on single-source data. To evaluate the benefits of multimodal integration, we used two evaluation strategies. The first strategy involved comparing our model with methods focused on unimodal clustering. In earlier sections, we compared scCODI with existing single-cell clustering methods (SC3 and TSCAN) and two general clustering methods (IDEC and K-means), which are commonly used for scRNA-seq data. We extracted RNA gene expression matrices, chromatin accessibility matrices, and protein expression matrices from the four previously mentioned RNA and ATAC multimodal datasets and from five CITE-seq datasets. These matrices were then subjected to cluster analysis, and their performance was evaluated using NMI and ARI. The experimental results demonstrate the advantages of multimodal integration over unimodal methods (Figure 3a and Figure 3b).

Additionally, because scCODI’s encoder is capable of performing cluster analysis on single-modal data, we got three scCODI variant models by keeping one omic dataset while setting the other omic data to all-zero values: a sub-model of scCODI with only RNA input and reconstruction loss (referred to as scCODI-RNA), and sub-models with only ADT or ATAC input and reconstruction loss (referred to as scCODI-ADT and scCODI-ATAC, respectively). scCODI outperformed the three variant models consistently across CITE-seq, SMAGE-seq, and SNARE-seq datasets; scCODI-RNA achieved the second-best performance (Figure 5a). This result aligns with our expectations because scRNA-seq data provide richer gene expression information and higher feature dimensions, which allowed the model to learn and to predict cell states more comprehensively.

**Figure 5.**
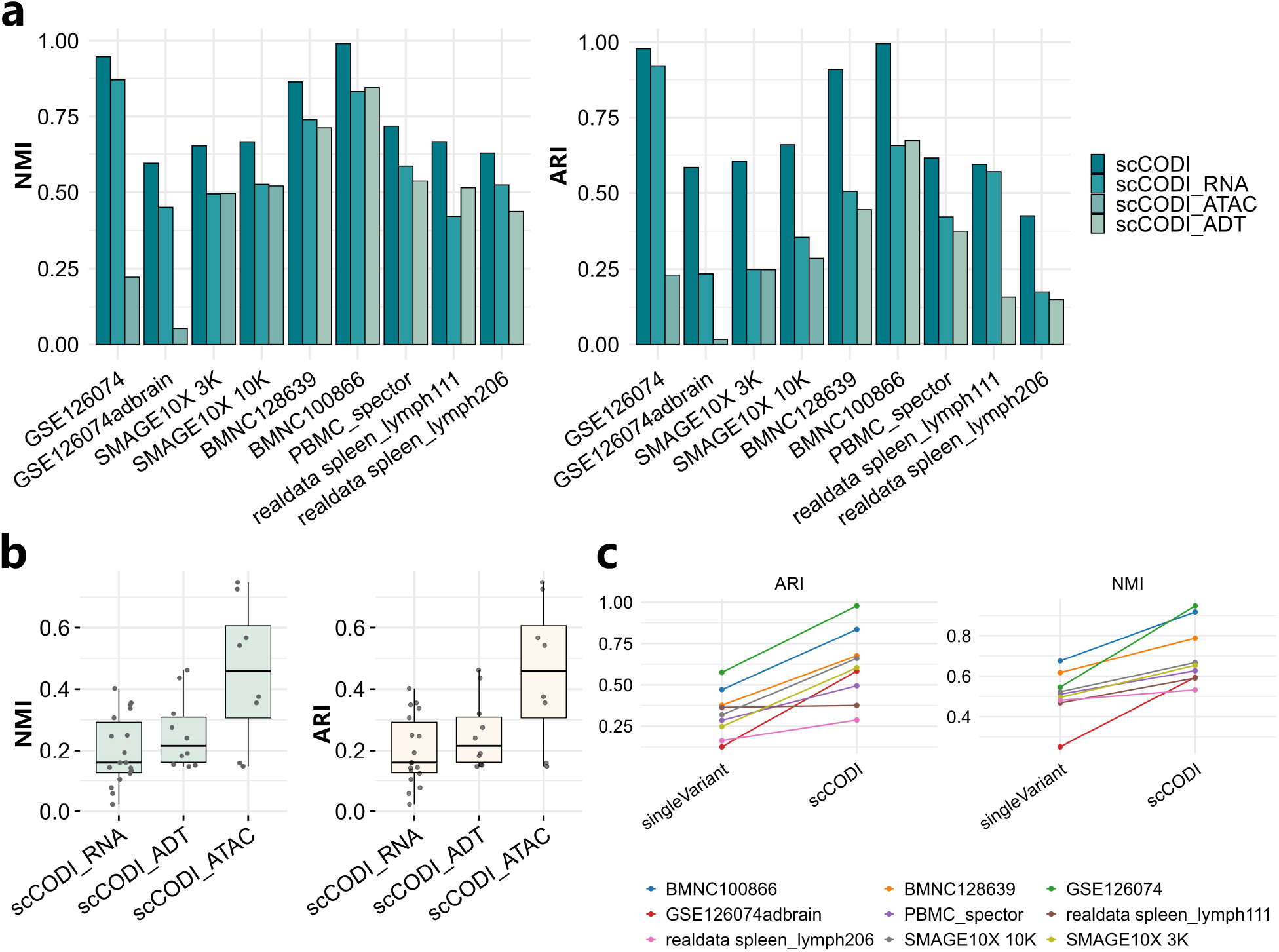
Clustering performance of scCODI and its variant models on multimodal datasets. scCODI, scCODI-RNA, and scCODI-ADT were evaluated across five CITE-seq datasets, and scCODI, scCODI-RNA, and scCODI-ATAC were evaluated across four SMAGE-seq datasets and SNARE-seq. (a) shows present clustering performance as a bar chart. (b) illustrates the performance differences between scCODI and other methods using box plots, separately for the CITE-seq datasets (left, n = 5) and the SMAGE-seq datasets (right, n = 4). Each box plot shows the minimum, first quartile (Q1), median, third quartile (Q3), and maximum values. The interquartile range (IQR), which is the difference between the third quartile (Q3) and the first quartile (Q1). The minimum and maximum are determined as the smallest and largest data points that are greater than or equal to Q1 - 1.5IQR and less than or equal to Q3 + 1.5IQR, respectively. The comparison between scCODI and its variants is shown in line plots in panel (c). Lines connect the paired performance for each dataset between the two methods.

In contrast, ADT data contained fewer features and more limited information, which resulted in lower performance when used alone compared with scRNA-seq data. On the other hand, scCODIATAC exhibited poorer performance across the four SNARE-seq and SMAGE-seq datasets (Figure 5b). Subsequently, we selected the higher-performing subset of the variant models across nine datasets for comparison with scCODI. There was a clear upward trend in performance with multimodal integration (Figure 5c).

scCODI is capable of integrating information from multimodal data to enhance clustering performance. To this end, we constructed UMAP plots (Figures S10) for scCODI and the three variant models.

### 3.5. Downstream analysis

To validate the clustering results derived from the latent representations, we conducted two standard downstream analyses: differential expression (DE) analysis and gene set enrichment analysis (GSEA) [40]. DE analysis was performed using the Scanpy package, which enables pairwise comparisons between two clusters or between a target cluster and the remaining clusters. Based on the ranked gene lists and their directions, GSEA was applied to identify enriched pathways within target clusters. As an example, we present the results from the BMNC100866 dataset (GSE100866), which focused on five clusters: B cells, CD14+ monocytes, CD4+ T cells, CD8+ T cells, and NK cells. DE and GSEA analyses were performed for each cluster compared to the others. The differentially expressed genes for these five cell clusters exhibited numerous well-known marker genes for each cell type (Figure 6). For example, LYZ was highly expressed in CD14+ monocytes [41]. MS4A1 is a marker gene for B cells, CD8A and CD8B are marker genes for CD8+ T cells [42], and NKG7 is highly expressed in NK cells [43].

**Figure 6.**
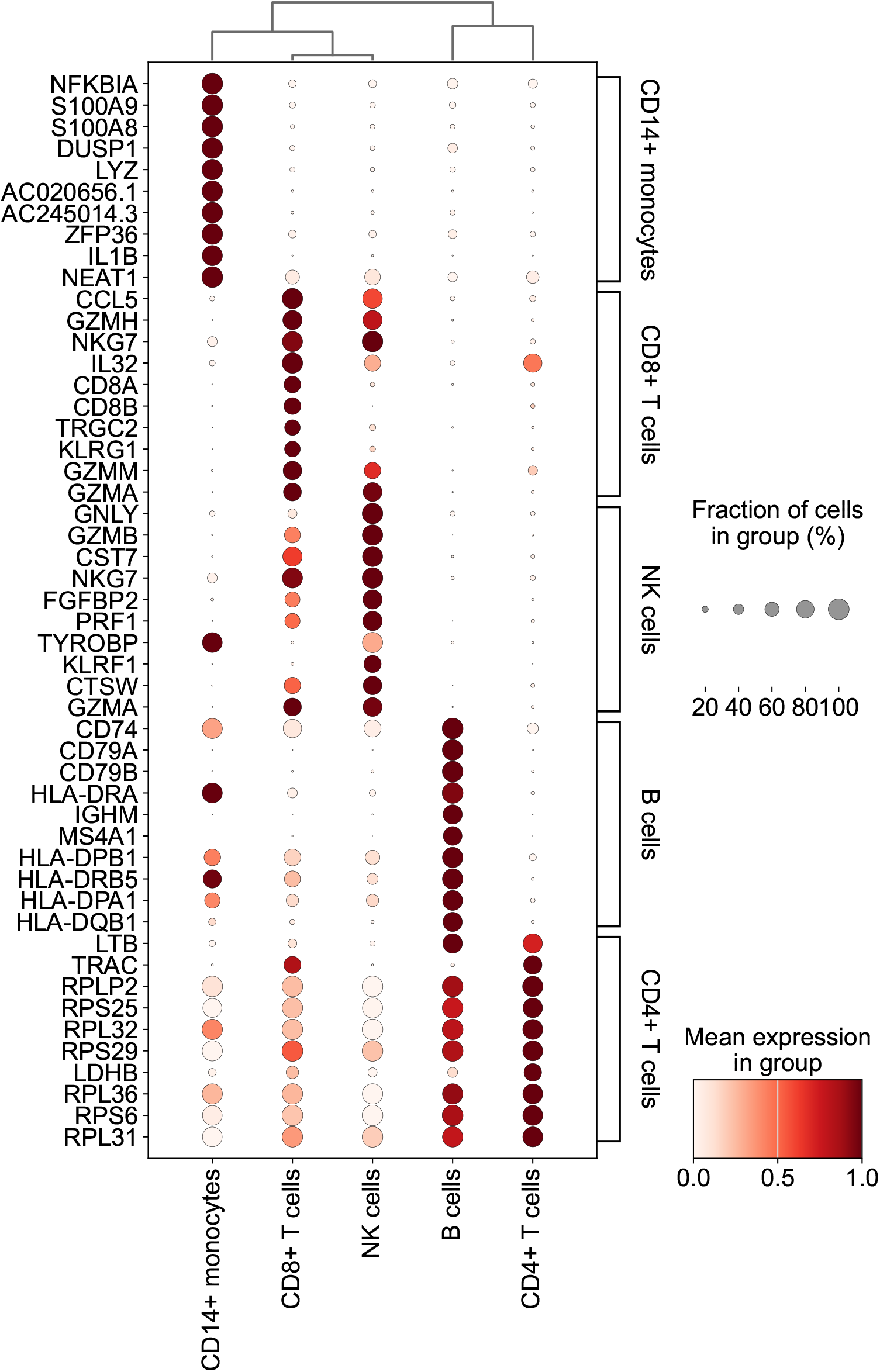
Dot plot visualization of marker gene expression across cell clusters. The dot plot illustrates the expression levels and the fraction of cells expressing specific marker genes across different cell types, including CD14+ monocytes, CD8+ T cells, NK cells, B cells, and CD4+ T cells. The size of the dots represents the percentage of cells in each group expressing a given gene, while the color intensity indicates the mean expression level of the gene within the group. Hierarchical clustering is applied to group genes and cell types based on their expression patterns.

The analysis of enriched pathways revealed the significant roles of different cell types in specific functional pathways (Figure 7). For instance, NK cells showed a marked enrichment in the “Natural killer cell-mediated cytotoxicity” pathway, highlighting their crucial role in immune defense [44, 45]. CD14+ monocytes exhibited significant enrichment in pathways such as “Antigen processing and presentation” and “NF-kappa B signaling pathway”, which further highlight their central role in immune regulation and inflammatory response [46].CD8+ T cells were notably enriched in the “T cell receptor signaling pathway”, which indicate the importance of this pathway in T cell function and antiviral defense [47, 48]. Similarly, CD4+ T cells were enriched in the “Antigen processing and presentation pathway”, which demonstrate their critical role in immune response and cellular activation [49, 50]. B cells showed significant enrichment in pathways, such as the “Intestinal immune network for IgA production” and the “Complement system”, which emphasizes their pivotal role in humoral immunity [51]. These findings further validate the specificity of functional activities across different immune cell types and support the accuracy and reliability of the clustering results obtained through our method.

**Figure 7.**
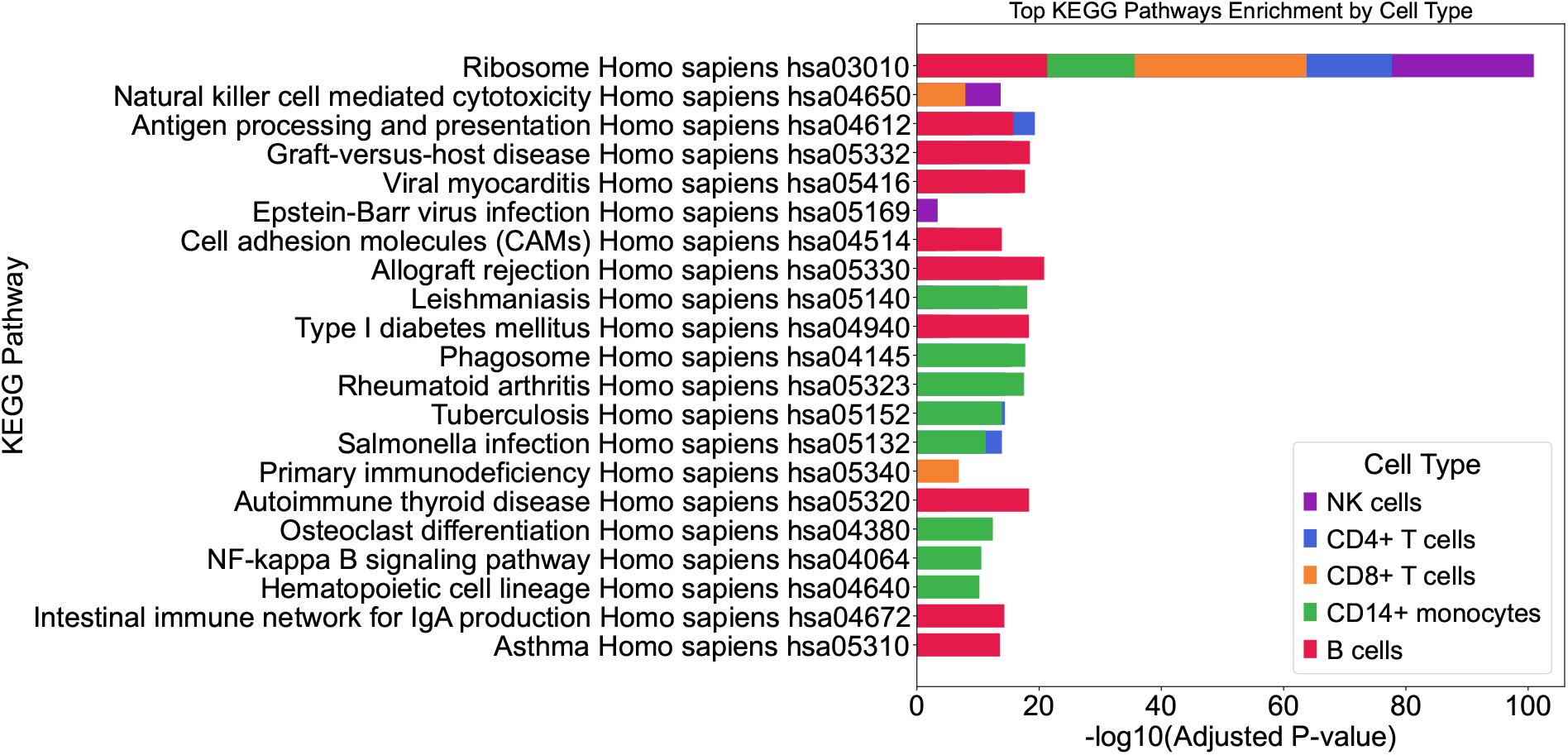
KEGG Pathway Enrichment Analysis by Cell clusters. This bar chart illustrates the KEGG pathway enrichment results for different cell types, including NK cells, CD4+ T cells, CD8+ T cells, CD14+ monocytes, and B cells. Each bar represents a KEGG pathway, with its length indicating the −log10 (adjusted P-value) as a measure of significance. Different colors correspond to different cell types. The pathway names are listed on the left, and longer bars indicate higher enrichment significance for the corresponding cell type in the respective pathway.

### 3.6. Ablation study and parameter sensitivity analysis

#### Convergence analysis

To verify the convergence of the model, we analyzed the iterative changes in the objective values and evaluation metrics (Figure 8). The results demonstrated that while the objective value generally decreased over iterations, minor fluctuations were observed during the process before it eventually stabilized. Similarly, the NMI and ARI values showed an initial gradual increase during the iterative process, followed by stabilization and minor fluctuations within a narrow range. These observations confirmed the convergence behavior of scCODI.

**Figure 8.**
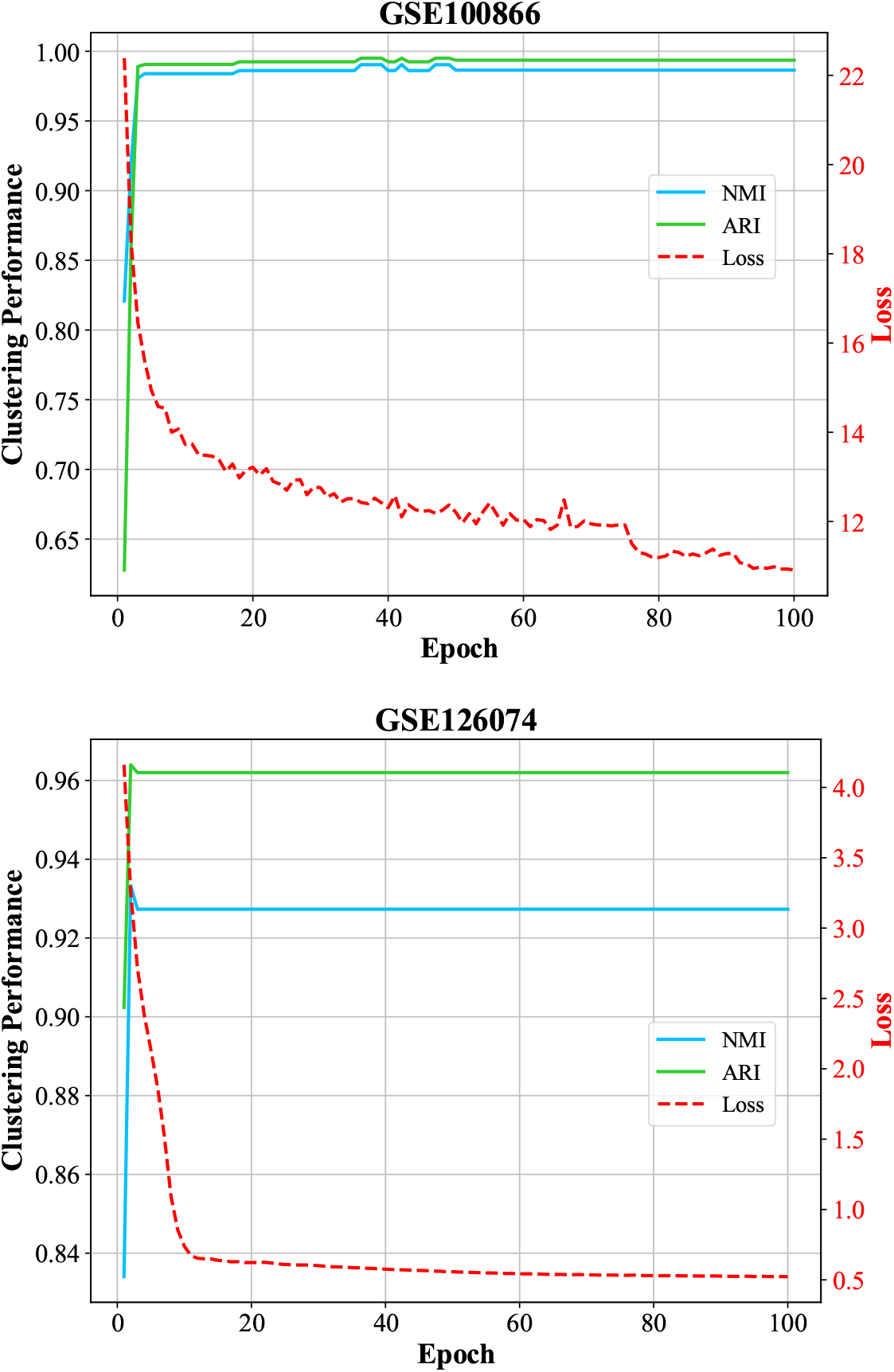
Clustering Performance and Loss Curve. This line plot illustrates the clustering performance and loss curve over training epochs for the BMNC100866 and GSE126074 datasets. The Y-axis on the left represents clustering performance metrics, including NMI and ARI, while the Y-axis on the right represents the loss value. The X-axis indicates the number of epochs. The blue and green lines represent the NMI and ARI scores, respectively, demonstrating convergence as training progresses. The red dashed line represents the loss, which decreases rapidly during the initial epochs and stabilizes as training continues.

#### Parameter sensitivity analysis

We evaluated the effect of hyperparameters on the clustering performance of scCODI, which included the trade-off coefficients *λ, μ, γ*, and the temperature parameter *τ*. NMI and ARI of scCODI when *λ* varied from 10^−2^ to 10^2^, *μ* varied from 0 to 1, *γ* varied from 10^−2^ to 10^2^, and *τ* varied from 0.2 to 0.8. The clustering results of scCODI were insensitive to *λ* and *τ* when *λ* was in the range of 0.1 to 1, when *μ* was in the range of 0.001 to 0.005, when *γ* is in the range of 0.01 to 0.1, and *τ* is in the range of 0.3 to 0.5 (Figure 9). Empirically, we set *λ, μ, γ*, and *τ* to 1.0, 0.001, 0.1, and 0.5, respectively.

**Figure 9.**
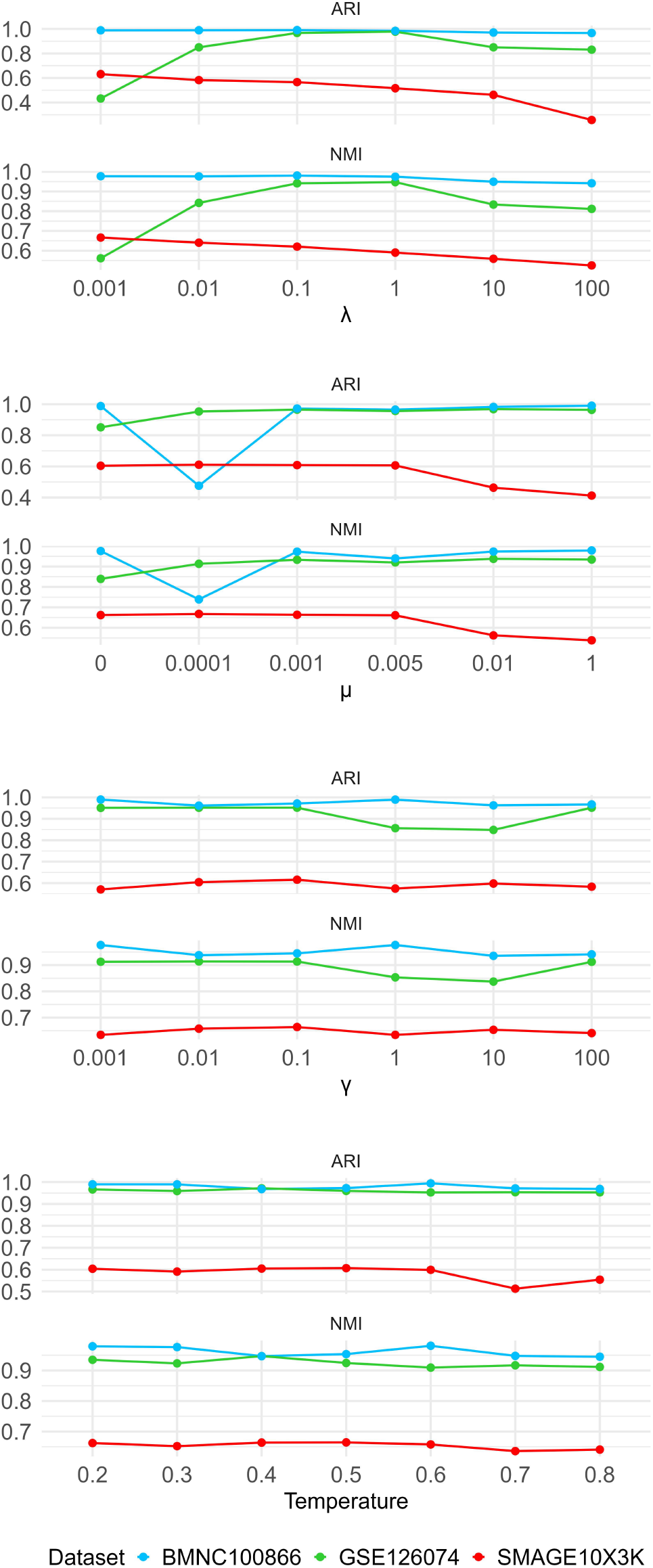
Hyperparameter tuning of scCODI. This figure demonstrates the impact of varying hyperparameters (*λ, μ, γ*, and temperature) on clustering performance metrics, including ARI and NMI.

#### Ablation study

We performed an ablation study to assess the contribution of each component in the proposed model. For this purpose, we implemented two alternative models and compared their performance with that of the proposed method.

##### Effectiveness of the COFInt module

Because the output of the COFInt module was the input to the GRgCL module, we set the resulting global relationship matrix as an identity matrix. Specifically, we used the low-dimensional embedding (*z*) obtained by concatenating the raw data of the two omics through an autoencoder and analyzing it in conjunction with the identity matrix for subsequent analysis. The results of the proposed method are higher in terms of NMI than the No-COFInt model by 9.87%, 9.94%, and 4.73% on the three datasets, respectively (Table 3). This indicated that the concatenated representation *Z* contained a large amount of view-specific information, which was not conducive to clustering. Although the No-COFInt model leveraged the complementarity of similar samples to reduce noise and redundancy across different views effectively, our COFInt module enhanced the consistency of data representation within the same cluster further.

**Table 3.** Ablation study. The best results are highlighted in bold font.

|  | noCOFInt |  | noGRgCL |  | scCODI |  |
| --- | --- | --- | --- | --- | --- | --- |
|  | NMI | ARI | NMI | ARI | NMI | ARI |
| BMNC100866 | 0.9014 | 0.8089 | 0.9299 | 0.9422 | **0.9904** | **0.9950** |
| GSE126074 | 0.8617 | 0.8616 | 0.8288 | 0.8315 | **0.9474** | **0.9781** |
| PBMC_spetor | 0.6856 | 0.4810 | 0.7120 | 0.4667 | **0.7180** | **0.6149** |

##### Effectiveness of the GRgCL loss

The results of the proposed method were higher in terms of NMI than the No-CL model by 9.54%, 14.31%, and 0.84%, respectively (Table 3). Because the multimodal shared feature representation was improved by incorporating the global relationships across all samples. Contrastive learning further optimized this by maximizing the similarity between the shared feature representation and the view-specific representation of the same sample, while simultaneously minimizing the similarity between representations of different samples that exhibit low relationships; this improved clustering performance.

## 4. Conclusion

In this paper, we proposed a novel cross-omics deep learning method called scCODI for cluster analysis of single-cell multi-omics data. Our method is capable of handling two types of omics data: singlecell transcriptomic data paired with epigenomic data, and single-cell transcriptomic data paired with proteomic data. We designed a cross-omics integration module that enhances the common representation by learning the global similarities between samples, which brings similar samples closer in space. Additionally, through a contrastive learning module, scCODI aligns the shared features and specific representations of omics data, which ensures that paired samples from different omics have similar specific representations. Our experimental results demonstrated that scCODI outperformed previous methods on diverse single-cell multi-omics datasets.

Overall, scCODI improves data clustering performance by combining a global and cross-view feature aggregation module with a multi-view contrastive learning module. Our method not only fully leverages the complementarity of different omics data, but also enhances the consistency of data representation through contrastive learning, which results in more accurate clustering. These results indicate that scCODI has great potential and broad application prospects in handling complex single-cell multi-omics data.

## 5. Code and Data availability

The code is available at https://github.com/SXUI-BDSI/scCODI All data underlying this article are available in Gene Expression Omnibus, at https://www.ncbi.nlm.nih.gov/geo/

## 6. Author Contributions

**Conceptualization**: Yingjie Guo,Yuhua Qian.

**Formal analysis**: Yingjie Guo, Zhen Liang.

**Funding acquisition**: Yingjie Guo, Zhen Liang.

**Investigation**: Chenyang Cheng,Yingjie Guo, Zhen Liang

**Methodology**: Yingjie Guo, Chenyang Cheng, Zhen Liang.

**Project administration**: Yingjie Guo, Zhen Liang,Yuhua Qian.

**Resources**: Yuhua Qian.

**Software**: Chenyang Cheng, Zheqing Zhu, Jin Zhou.

**Supervision**: Yingjie Guo, Zhen Liang,Yuhua Qian.

**Visualization**: Chenyang Cheng, Zheqing Zhu, Jin Zhou.

**Writing – original draft**: Chenyang Cheng, Yingjie Guo.

**Writing – review & editing**: Chenyang Cheng, Haiping Lv, Yingjie Guo, Zhen Liang, Yuhua Qian.

## 7. Funding

This research was supported by the Science and Technology Innovation Young Talent Team of Shanxi Province to ZL (NO. 202204051001019), the National Natural Science Foundation of China to ZL (NO. 32170410), the Natural Science Foundation of Shanxi Province for the Excellent Youth to ZL (NO. 202203021224002), Fundamental Research Program of Shanxi Province to YG (NO. 202403021221020), Shanxi Scholarship Council of China to YG (NO. 2023-006). The funders had no role in study design, data collection and analysis, decision to publish, or preparation of the manuscript.

## Notes

### Competing Interest Statement

The authors have declared no competing interest.

