## Supplementary Figure S1-S10 for "Global and cross-omics feature aggregation improves single-cell multi-omics integration and clustering"

April 10, 2025

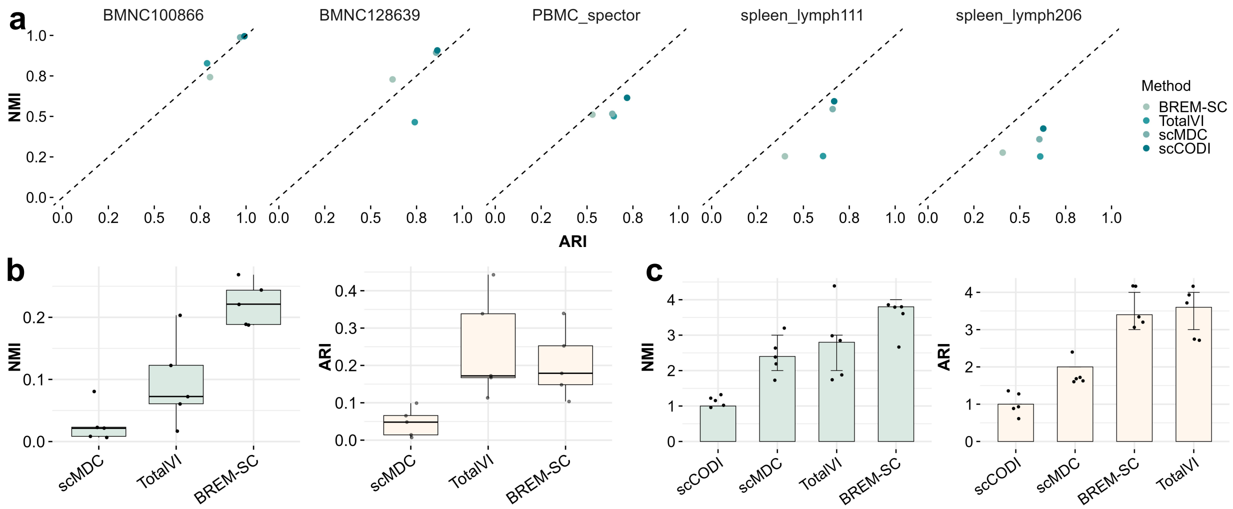

Figure S1: The clustering performance of scCODI and competing methods on various CITE-seq datasets. All methods were tested on five datasets. Panels (a) of the chart present clustering performance in a two-dimensional manner. Panel (b) illustrates the performance differences between scCODI and other methods using box plots, which display statistical summaries that include the minimum, first quartile (Q1), median, third quartile (Q3), and maximum values. The minimum represents the smallest data point equal to or greater than Q1 - 1.5IQR, and the maximum represents the largest data point equal to or less than Q3 + 1.5IQR. Each data point (representing a performance difference in a dataset) is shown by a dot. Additionally, panel (c) provides an overview of the average ranking for each method, with dots indicating the rank of a method for a specific dataset and error bars showing standard errors.

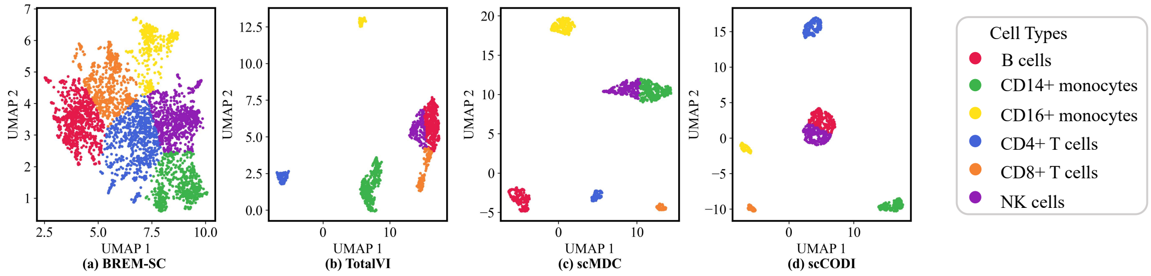

Figure S2: Cell clustering visualization of each method on GSE100866. (a) BREM-SC, (b) TotalVI, (c) scMDC, and (d) scCODI.

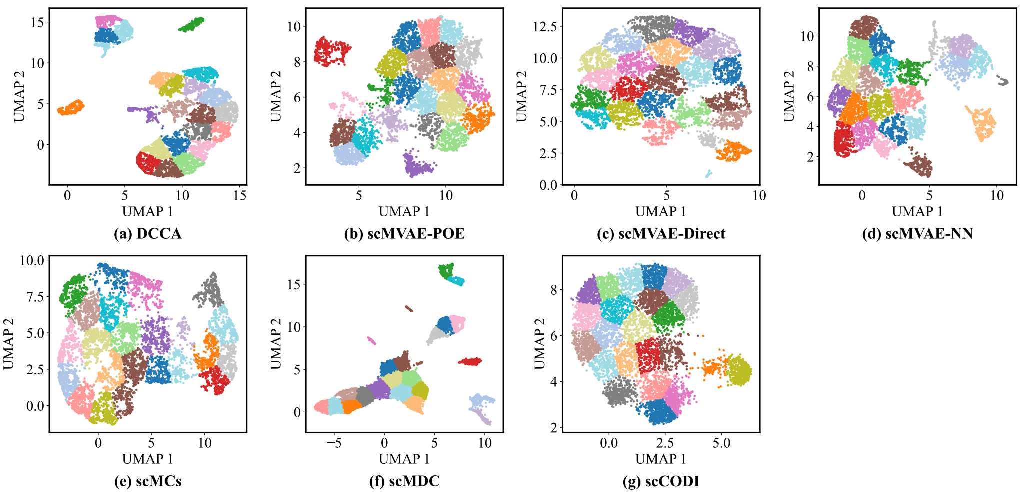

Figure S3: Clustering visualization for AdBrain. (a) DCCA, (b) scMAVE-POE, (c) scMVAE-Direct, (d) scMVAE-NN, (e) scMCs, (f) scMDC, (g) scCODI.

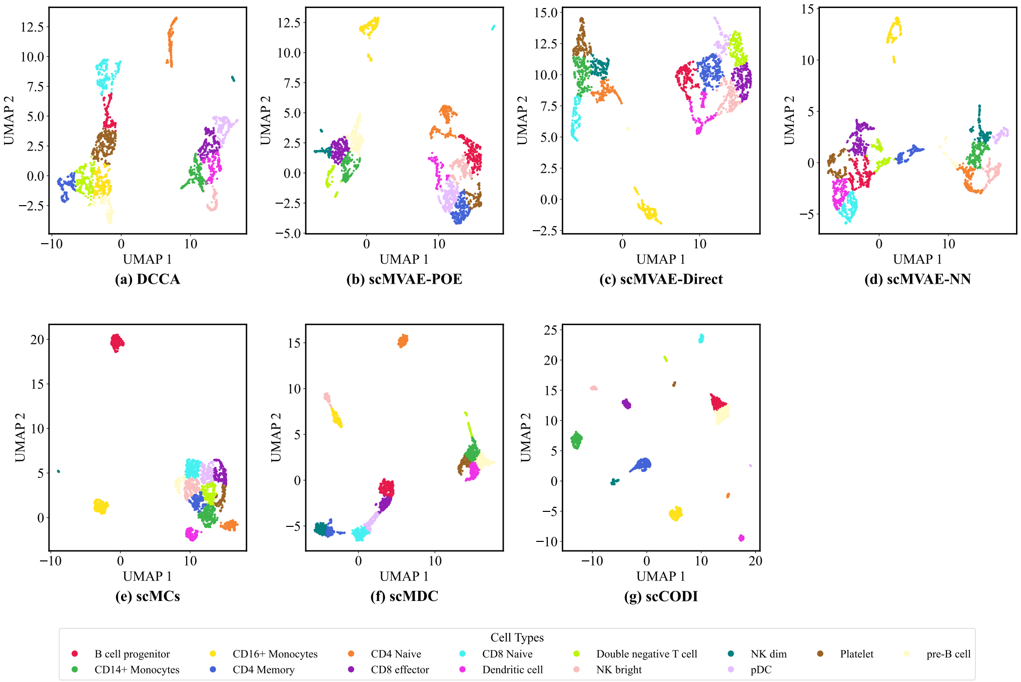

Figure S4: Clustering visualization for SMAGE10X 3K. (a) DCCA, (b) scMAVE-POE, (c) scMVAE-Direct, (d) scMVAE-NN, (e) scMCs, (f) scMDC, (g) scCODI.

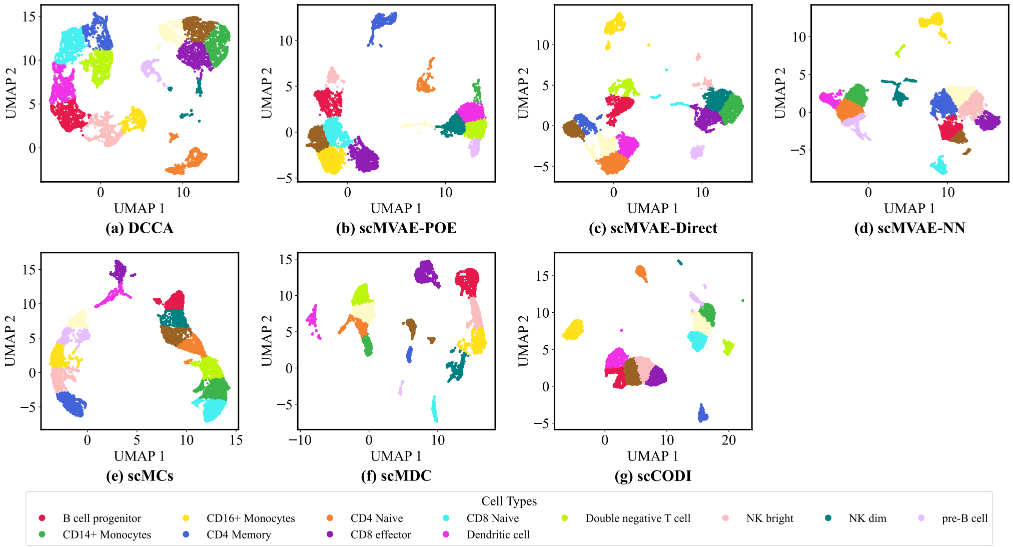

Figure S5: Clustering visualization for SMAGE10X 10K. (a) DCCA, (b) scMAVE-POE, (c) scMVAE-Direct, (d) scMVAE-NN, (e) scMCs, (f) scMDC, (g) scCODI.

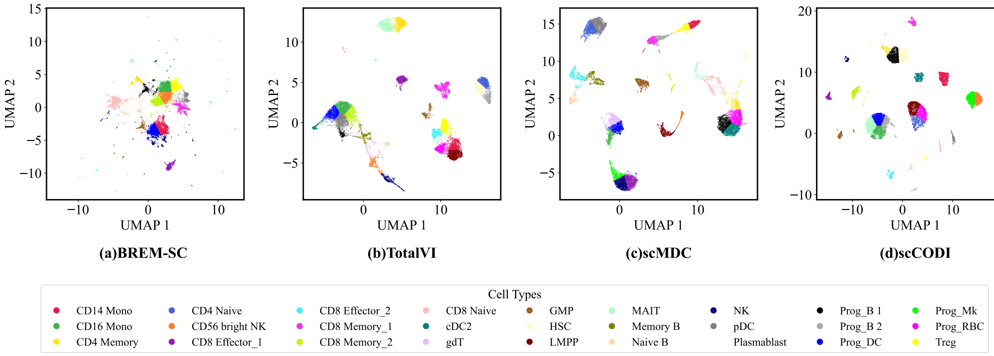

Figure S6: Clustering visualization for BMNC128639. (a) BREM-SC, (b) TotalVI, (c) scMDC, (d) scCODI.

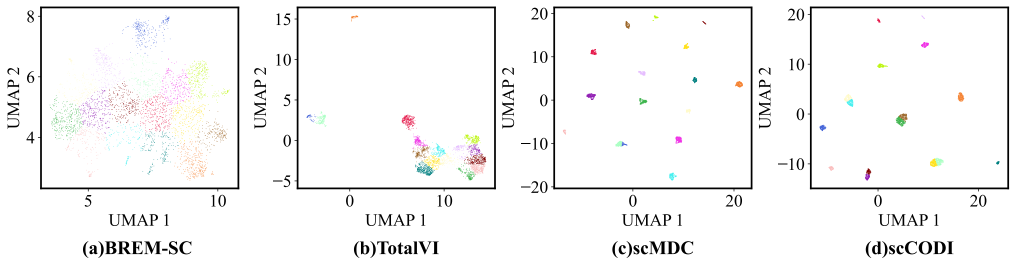

Figure S7: Clustering visualization for spector. (a) BREM-SC, (b) TotalVI, (c) scMDC, (d) scCODI.

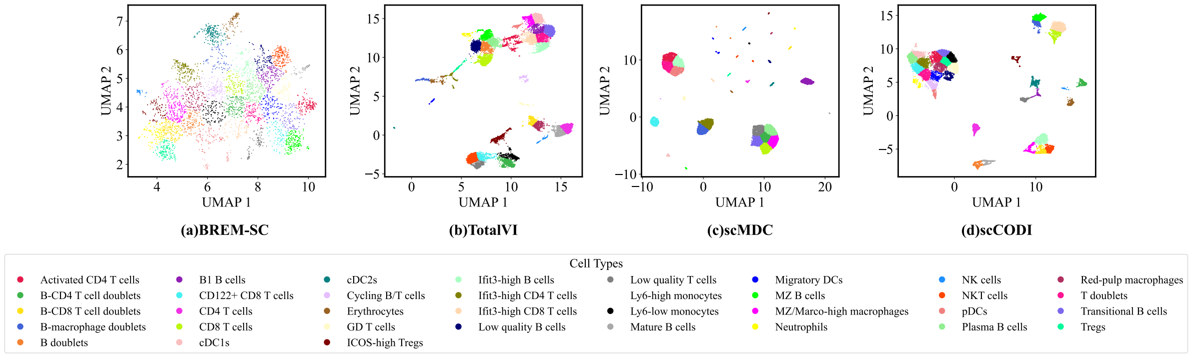

Figure S8: Clustering visualization for sln111. (a) BREM-SC, (b) TotalVI, (c) scMDC, (d) scCODI.

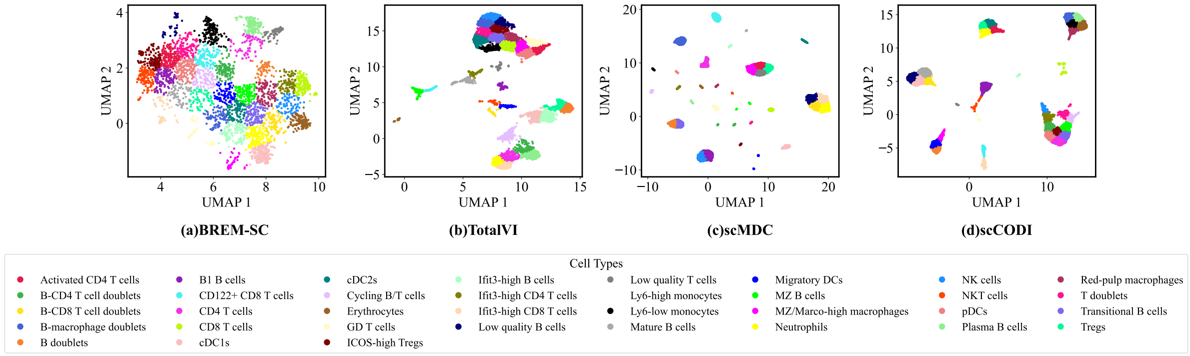

Figure S9: Clustering visualization for sln206. (a) BREM-SC, (b) TotalVI, (c) scMDC, (d) scCODI.

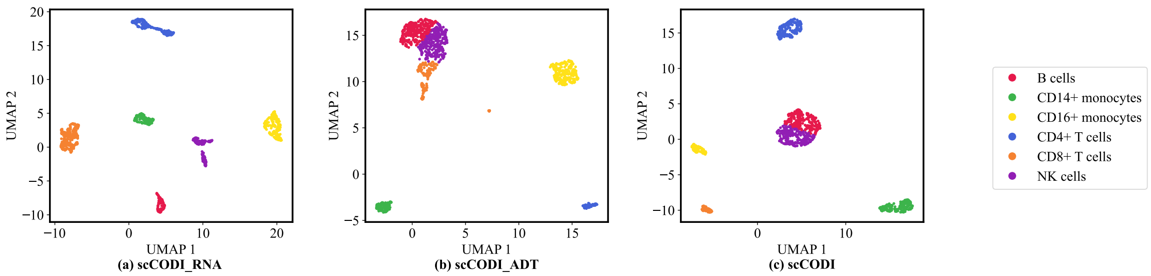

Figure S10: The clustering results of scCODI on the GSE100866 dataset across three variant models. (a) scCODI-RNA, (b) scCODI-ADT, (c) scCODI.
